# Motor decision-making under uncertainty and time pressure

**DOI:** 10.1101/2024.09.02.610909

**Authors:** Samuele Contemori, Timothy J. Carroll

## Abstract

Purposeful movement often requires selection of a particular action from a range of alternatives, but how does the brain represent potential actions so that they can be compared for selection, and how are motor commands generated if movement is initiated before the final goal is identified? According to one hypothesis, the brain averages partially prepared motor plans to generate movement when there is goal uncertainty. This is consistent with the idea that motor decision making unfolds through competition between internal representations of alternative actions. An alternative hypothesis holds that only one movement, which is optimised for task performance, is prepared for execution at any time. Under this conception, decisions about the best motor goal given current information are completed upstream from neural circuits that perform motor planning. To distinguish between these hypotheses, we modified Alhussein and Smith (2021) experiment in which participants had to start reaching toward targets associated with opposite curl force-fields prior to knowing the correct target to reach. Crucially, we forced the participants to initiate movement immediately after target presentation (i.e. mean reaction times ∼250ms) so that they had limited opportunity to deliberate between the available alternatives. We found that the reaching dynamics reflected only those learnt for the selected reach direction, rather than a combination of those for the alternative targets presented, irrespective of the time available to initiate movement. The data are consistent with the conclusion that reaching dynamics were specified downstream of action selection under the target uncertainty conditions of this study.

**NEW & NOTEWORTHY:** Here we found no evidence of “motor averaging” of reach dynamics for multiple potential actions when people had to respond as quickly as possible to uncertain target location cues. People exerted forces appropriate for the specific reach direction they selected irrespective of movement initiation time, suggesting that reaching dynamics were specified downstream of action selection.

## INTRODUCTION

Most movements that animals make in natural settings require selection of a particular action from a range of alternatives; with some uncertainty over which action will best meet the animal’s needs. This implies that multiple actions should be represented in some form within the brain so that they can be compared for selection. However, there remains controversy about the nature of the neural dynamics that underlie motor decision making, and the algorithmic principles that define motor behaviour under action uncertainty.

Much of this controversy has focussed on the outcomes of laboratory experiments that employed the so called “go before you know” paradigm (e.g. Chapman et al., 2010; McPeek et al., 2003; Watanabe and Higuchi, 2022; see Gallivan et al., 2018 for review). Here, behaviour and/or neural activity are studied when a movement is required before the task goal is specified from multiple alternatives. In perhaps the simplest and most common form of this task, two potential visual targets for reaching are presented simultaneously. Only after the reach is initiated is the correct end point identified. Thus, participants must plan and initiate movement before knowing which of the alternatives will ultimately be specified as the desired action. Classical behaviour in this task is for participants to launch their reaches at an intermediate angle between the two target alternatives (e.g. Stewart et al., 2013, 2014; Gallivan et al., 2011, 2015, 2016, 2017; Gallivan and Chapman, 2014; Van der Stigchel et al., 2006; Enachescu et al., 2021; Mangin et al., 2023), although direct reaching to one or the other of the targets can also be observed depending on task constraints (Haith et al., 2015a; Wong and Haith, 2017; Wong et al., 2022).

Many interesting and clever variants of this basic “go before you know” paradigm have been tested to dissociate sensory from motor effects, and to manipulate task constrains (e.g. Stewart et al., 2013, 2014; Gallivan et al., 2011, 2015, 2016, 2017; Gallivan and Chapman, 2014; Van der Stigchel et al., 2006; Alhussein and Smith, 2021; Enachescu et al., 2021; Mangin et al., 2023; Haith et al., 2015a; Wong and Haith, 2017; Wong et al., 2022). At the behavioural level, the results have been interpreted according to a theoretical dichotomy. On the one hand, observations of intermediate reaching behaviour are often interpreted as evidence that the brain averages “partially-prepared motor plans” to generate movement when uncertainty about desired action is unresolved (Cisek and Kalaska, 2005; Cisek, 2007; Chapman et al., 2010; Stewart et al., 2013, 2014; Gallivan et al., 2011, 2015, 2016, 2017 Gallivan and Chapman, 2014; Van der Stigchel et al., 2006; Enachescu et al., 2021; Mangin et al., 2023). This is the so-called “motor encoding” (e.g. Stewart et al., 2014; Cisek & Kalaska, 2010; Enachescu et al., 2021) hypothesis, which is aligned with the more general theory that decision making is achieved through competition between internal representations of alternative actions (Cisek, 2007; Cisek and Kalaska, 2010; Hunt and Hayden, 2017). On the other hand, observations of direct reaching to one of the alternative targets, or motor behaviour that cannot be explained as a weighted average of the two alternative actions, has been interpreted as evidence that the brain only ever plans a single movement that is optimised for task performance (Haith et al., 2015a; Wong and Haith, 2017; Wong et al., 2022). This is the so-called “performance-optimisation” hypothesis. In this conception, decisions about the best motor goal given current information and constraints are completed upstream from networks that determine how the motor goal will be achieved (i.e. before “motor planning”; Wong et al, 2014), so that only a single movement is prepared for execution at any time.

This theoretical dichotomy to explain motor behaviour under uncertainty has parallels with debates about neural correlates of motor decision making. If multiple alternatives for action are to be considered by the brain, there must be competing neural activity that represents those actions in some form. At one end of the theoretical spectrum, competition between actions could occur in parallel throughout all brain areas that contribute to sensorimotor behaviour. This idea is encapsulated by Cisek’s “affordance competition hypothesis”, which rejects the traditional notion of a serial transition from perception to decision-making and then action, and views brain activity as a set of parallel processes designed to identify and select between potential motor actions (Cisek, 2007; Cisek and Kalaska, 2010). In support of this idea, there is an extensive literature showing that multiple cortical regions represent multiple potential targets or actions under uncertainty (see Cisek & Kalaska 2010; Hunt & Hayden, 2017; Gallivan et al. 2018 for review). However, controversy remains over the extent to which activity associated directly with preparation and execution of movement incorporates information about multiple alternatives for action. A key consideration here is that classical, single-electrode, extracellular brain recording approaches require averaging across many trials and sessions to provide insight about movement representations, which can result in misleading conclusions under some circumstances. For example, Dekleva and colleagues (2018) re-examined classical evidence obtained from single-electrode recordings that dorsal premotor cortex represents two potential reach directions (Cisek and Kalaska, 2005). By simultaneously recording multiple neurons, the authors showed motor cortical dynamics consistent with preparation of movement to only one target at a time.

If we are to resolve these controversies, it seems clear that close attention to the task details that give rise to behaviour and neural activity must be paramount. In general, primates are capable of considerable flexibility in how they respond to stimuli depending on the behavioural context, so caution is required when attempting to extrapolate findings obtained in a specific context to universal principles. One key task constraint is the time available to process the stimuli that define action alternatives before movement is required. Given sufficient time, primates can produce a large variety of movements with flexible relationships to the physical characteristics of target stimuli. For example, humans can readily look or reach away from the target direction in anti-saccade (e.g. Everling & Fischer, 1998; Munoz & Everling, 2004), anti-reach (e.g. Connolly et al., 2000, Gu et al., 2016), or target-jump tasks (e.g. Leow et al, 2017; 2020), but at the expense of longer reaction times. Thus, any viable “motor encoding” theory must include scope for internal representations of the desirability of actions to contribute to movement specification, as has been proposed (Cisek, 2007; Enachescu et al 2021). Moreover, the longer the time available to process stimuli that define action alternatives prior to movement release, the greater the opportunity for any competition between brain representations of these alternatives to be resolved. The implication is that competition or averaging of “motor plans” is more likely to be evident soon after target stimuli are presented than after an extended duration.

Most of the behavioural studies cited to support the idea that optimization principles drive behaviour in “go before you know” tasks allowed participants ample time (>1s) to view the potential targets before movement initiation (Haith et al., 2015a; Wong and Haith, 2017; Alhussein and Smith, 2021; Wong et al., 2022). In particular, Alhussein and Smith (2021) recently reported elegant and influential experiments that were argued to provide strong evidence in favour of the performance-optimization theory. In one of these experiments, people adapted to force-fields associated with three potential reach targets. The force field for the two lateral targets had the opposite sign to the force field for the central target, such that different reach dynamics were required to reach to different parts of the workspace. When the two lateral targets were presented as potential targets in a “go before you know” condition, participants’ reach dynamics were consistent with a single motor plan encoding the central target direction, rather than an average of the motor plans for the two lateral targets. Although the results are clearly consistent with a performance optimisation interpretation, because participants were given considerable time (>1s) after target presentation to initiate a response, they were not challenged to move prior to an opportunity to deliberately choose among the available alternatives. Until this is addressed, definitive conclusions about the default sensorimotor transformation mechanisms in circumstances of action uncertainty cannot be drawn.

Here, we searched evidence of “motor averaging” of reach dynamics for alternative targets in a task context more likely to reflect such behaviour. We replicated the paradigm designed by Alhussein and Smith (2021) but forced the participants to initiate movement immediately after target presentation. We applied a hard time limit of 500ms to initiate reaching, which resulted in mean reaction times of ∼250ms. We perturbed the reaching trajectory toward straight-ahead (central) and lateral (right\left) targets via opposite curl force-fields. We then tested which of the newly learnt reaching dynamics was adopted when the lateral targets appeared simultaneously in a “go before you know” condition. If motor averaging occurs at the level of reach dynamics specification, we predicted that participants should exert forces consistent with the lateral target force fields for the shortest reaction time trials. By contrast, if reaching dynamics are specified downstream of action selection, participants should exert forces that reflect only the dynamics learnt for central target irrespective of the time available to initiate their response after stimulus presentation.

## MATERIALS AND METHODS

### Participants

Twenty-four healthy right-handed adults (8 females; mean age: 27±7 years) participated in this study. All participants had normal or corrected-to-normal vision and no neurological and\or musculoskeletal disorder at the time of the experiment. They provided informed consent, received payment ($20/h) for their participation, and were free to withdraw from the experiment at any time. All experimental procedures were approved by the University of Queensland Medical Research Ethics Committee (Brisbane, Australia) and conformed to the Declaration of Helsinki.

### Experimental set-up and trial structure

The participants performed visually-guided reaches using a two-dimensional planar robotic arm (vBOT, Figure 1A; Howard and Ingram, 2009). The task paradigm was created in Microsoft Visual C++ (Version 14.0, Microsoft Visual Studio 2005) using the Graphic toolbox. The participants received visual feedback about the stimuli and hand movement via a mirror that reflected an LCD computer monitor (120Hz refresh rate) and occluded direct vision of the arm (Figure 1A). The hand position was virtually represented by a red cursor (1cm in diameter; Figures 1A-D) whose apparent position coincided with the actual hand position in the plane of the limb (Figure 1A).

**Figure 1:**
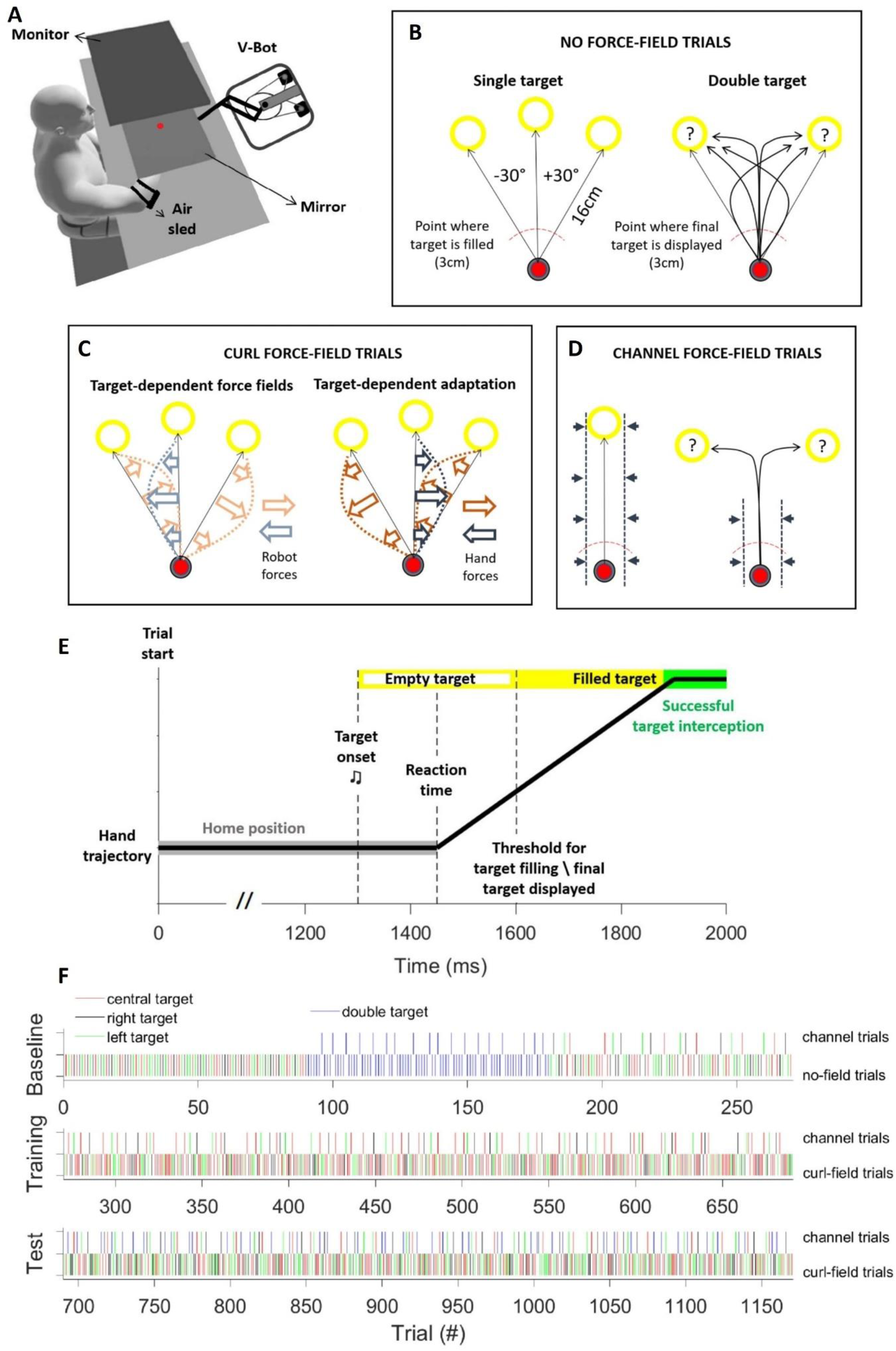
A, Experimental setup. The head position was stabilized by a forehead rest (not shown here) and the upper arm was supported on a custom-built air sled positioned under the right elbow to minimize arm-on-table sliding friction. B, Schematic representation of the no force-field trial types. In the single target trials, the target could appear at 0° (central target), +30° (right target) and −30° (left target) relative to the body midline. In the double-target trials, the right and left targets were presented simultaneously (‘?’ = uncertainty about the final reaching goal) until one of them was filled and the other extinguished when the hand crossed the movement threshold (red dashed arc). C, Schematic representation of the curl force-field trials wherein the robot applied opposite target-dependent and velocity-dependent forces on participants’ reaching hand. In this example, the robot applied counter-clockwise forces for central targets and clockwise forces for lateral targets, thus requiring opposite adaptive forces applied on the robot handle. D, The hand-on-handle forces were measured as the force required by the robot to constrain the hand trajectory within a virtual channel. In the single target trials, the channel was applied along the entire target-directed direction (note that only the central target is shown for simplicity). In the double-target trials, the channel was oriented between the two possible targets, but was applied only up to half of the home-to-target distance (i.e. 8cm). E, Timeline of events for a single trial. The cursor had to stay at the home position for 1300ms after which a yellow ring-shaped target was presented simultaneously with a chirp tone to cue the necessity of a motor response. The target was filled once the hand crossed a virtual movement threshold (i.e. 3cm from the home position; see panel B). In the double-target conditions, this event corresponded to the point in time at which the final target was displayed, and the distractor target turned off. F, Testing protocol and exemplar distribution of the different trial types across the three experimental phases. See Materials and Methods for further details.

To start a trial, the participants had to bring the cursor to a starting ‘home’ position (grey ring of 1.5cm in diameter; Figures 1B-D) aligned to the mid-body line and to hold this position for 1.3s by keeping the hand velocity <5cm/s (Figure 1E). In the single-target conditions, a yellow ring target of 2cm in diameter was then presented at 16cm from the home position simultaneously with an auditory signal (i.e. ‘go’ cue; Figure 1E). The ring target could appear at −30° (left target), 0° (central target), or +30° (right target) from the mid-body line (Figure 1B) and was filled as soon as the hand position was >3cm from the home position (i.e. target fill event; dashed red line in Figures 1B and D). In the double-target conditions, the right and left ring targets appeared simultaneously, and the participants had to start moving before knowing the final target to reach. At the target fill event (see above), the final target for reaching was filled and the distractor turned off (Figure 1B).

To avoid anticipatory responses, a “Too Early” error message was shown if the participants left the home position before the target presentation. To force early response initiation, a “Too Late” error message was shown if the hand left the home position >500ms after the target presentation. Pilot experiments showed that this reaction-time constraint allowed early movement initiation without substantial degradation in task performance. Consistent with the Alhussein and Smith (2021) study, the participants received “Too Fast”, or “Do Not Decelerate”, error messages if they moved faster than 8m/s, or reduced their instantaneous maximum velocity by more than 33%, during the first half of the movement (i.e. hand-from-home distance <8cm). This allowed us to preserve consistent movement parameters across conditions and testing phases, as well as avoid ‘exploratory’ behaviour in double-target trials (see Alhussein and Smith 2021). Any of the above error messages led to the immediate interruption of the trial, which was repeated at the end of the block.

To successfully complete the trial, the participants had to reach the target and remain within the target space by keeping the hand velocity <5cm/s for 300ms. At the end of the trial, the target was turned green (Figure 1E) if the reaching action was completed within 700ms or 1000ms from the movement onset for the single-target and double-target conditions, respectively. Otherwise, the target was turned blue simultaneously with the presentation of a “Too Slow” warning message.

### Curl and channel force-fields

To test how the brain represents multiple options for actions, we dissociated the reaching dynamics across the central and lateral target locations. Specifically, we perturbed the hand trajectory to the central and the lateral targets via opposite curl force-fields to induce distinct target-dependent adaptations (Figure 1C). The force-fields were generated by the vBOT motors and transmitted by the robot handle to the participants’ reaching hand, thus perturbing the target-directed trajectory. The forces were proportional to the hand velocity and orthogonal to the hand-velocity vector as described by the following equation:

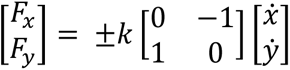

where k was the force gain (±25 N·s/m) whose sign determined the direction of the force field (+ clockwise CW; - counterclockwise CCW). Half of the participants experienced the CCW and CW curl force-fields for the central and lateral targets, respectively, whereas the other half experienced the opposite combination of target-dependent curl force-fields.

The degree of adaptation was evaluated by measuring the force applied by the hand on the robot handle when the movement was confined within a virtual mechanical channel that was generated by the vBOT motors by simulating a stiff spring and damper (10000 N/m, 100 N/m/s; Figure 1D; Carroll et al. 2019). Specifically, we calculated the hand force as the inverse of the robot force needed to constrain the hand trajectory within the virtual channel.

For the single-target trials, the channel was oriented toward the target and confined the hand trajectory along the whole reaching distance (Figure 1D). By contrast, for the double-target trials the channel was oriented at 0° relative to the body midline (i.e. between the two potential targets) and confined the hand trajectory for only half (8cm) of the target distance (Figure 1D). This allowed us to record the reaching dynamics prior to displaying the final target. Note that the robot-force gradually turned off the robot force over 100ms after the hand position reached 8cm from the home position, thus avoiding avoid reflexive responses to sudden state changes.

### Testing protocol

We replicated the testing protocol described in Alhussein and Smith (2021) as it proved effective to induce large target-dependent adaptation among all participants, despite the use of opposite target-dependent perturbing force-fields. The experiment consisted of three phases: (i) baseline; (ii) training; (iii) test (Figure 1F).

The baseline phase comprised two familiarization blocks of 45 single-target trials (15 trials * 3 targets * block). Subsequently, we evaluated participants’ reaching behaviour in the double-target conditions in three blocks of 30 trials/block composed by 24 no-field double-target trials (12 trials * 2 targets * block), which were randomly intermixed with 6 channel-field double-target trials (3 trials * 2 targets * block). Finally, the participants completed two blocks of 45 trials/block composed by a mixture of 36 single-target trials (12 trials * 3 targets * block) and 9 channel-field single-target trials (3 trials * 3 targets * block). The channel-field trials allowed us to obtain the reaching dynamics for the different target conditions prior to experiencing the perturbing force-fields.

During the training phase, the participants completed seven blocks of 60 single-target trials per block, with each block comprising 48 curl force-field trials randomly intermixed with 12 channel-field trials. Note that the central target was presented twice as frequently as the lateral targets to ensure balanced training across the opposite curl force-fields.

The test phase goal was twofold, as we wanted both to preserve the target-dependent adaptation acquired during the training phase and evaluate the newly learnt reaching dynamics in the double-target conditions. To these aims, the participants completed six blocks of 80 trials/block composed of 60 force-field trials in the single-target conditions (20 trials * 3 targets * block) randomly intermixed with 20 channel-field trials distributed among single (12 trials: 4 trials * 3 targets * block) and double (8 trials: 4 trials * 2 targets * block) target conditions.

Importantly, the channel trials were pseudo-randomly interspersed with the other trial types to avoid consecutive presentations of channel-field trials and decay in adaptation (Figure 1F).

### Data recording and analysis

The kinematic and dynamic data were recorded at 1 KHz via the vBOT optical encoders. The movement reaction time (RT) was defined via a technique previously (Veerman et al., 2008; Wijdenes et al., 2014; Zhang et al. 2018a, 2018b; Contemori et al., 2023). Briefly, we identified the first peak of the radial hand-velocity exceeding the baseline value (computed across the 100ms preceding the target onset) by >5 SDs. We then fitted a line to the hand velocity data enclosed between 25% and 75% of this peak and indexed the RT as the time of intersection between this line and the baseline velocity value. Trials in which the RT was <130ms (∼2%) were excluded during off-line analysis because 130ms was shown to be the minimum time needed to accurately prepare a purposeful target-directed response (Haith et al., 2016). We then defined the initial reach direction by taking the slope of a line connecting the hand-position coordinates at the RT and when it reached 2.5cm from the home position; that is prior to the target fill event (i.e. hand position >3cm from the home position; Figures 1B and E).

We also tested whether the target-goal uncertainty modulated the movement time. It is worth noting that the double-target conditions required a trajectory correction to intercept the final target, especially during the channel-field trials (Figures 1B and D). This was likely to increase the movement time relative to the single-target conditions even if the reach evolved at a similar speed across the conditions. Therefore, for all conditions we evaluated the movement time by taking the time difference between the time when the hand trajectory exceeded 8cm and the RT (i.e. half-movement time).

To measure adaptation to the perturbing force-fields, we first computed the hand-force from 300ms before to 300ms after the target fill event for each channel-field trial of the baseline phase. For each subject, we then pooled the baseline hand-force traces for each of the three single-target locations and calculated the average across trials. For the double-target conditions, the baseline hand-force was also averaged across the two (right and left) target locations. We then subtracted the baseline hand-forces from those recorded in compatible target-dependent conditions of the training and test phases. Subsequently, the hand-forces were normalized to the maximum value of the “ideal” force that, based on the actual hand velocity, would fully compensate the perturbing force-field up to 100ms after the target fill event. Importantly, this procedure prevented the velocity trace from being contaminated by the earliest encoding of the final target in the double-target conditions. In fact, ∼100ms is the time for both the earliest muscle (Kozak et al., 2019; Cross et al., 2019), postural (Fautrelle et al., 2010; Zhang et al., 2018a, 2018b), and movement (Day and Lyon, 2000; Day and Brown, 2001; Veerman et al., 2008; Smeets et al., 2016; Brenner et al., 2022) adjustments during ongoing reaching actions. This procedure returned normalized force-traces in which positive values denoted hand-forces directed congruently with the ideal compensatory force. Similarly, the normalized hand-force in the double-target condition was defined as positive when appropriate to compensate for the central target force-fields and negative when appropriate to compensate for the lateral target force-fields. Finally, we quantified adaptation by taking the normalized hand-force at the peak velocity.

To test the time-course of motor learning and retention, we averaged the adaptation metric for the first and second halves of the training phase, and for all test phase trials. For the lateral-target conditions, we also averaged the adaptation value across the right and left targets. For each participant, therefore, we obtained average adaptation values for central versus lateral target presentations in the early-training, late-training, and test phases.

In the double-target channel field conditions, participants sporadically (∼8% of relevant trials) produced rapid and extremely large forces that were up to 4 times larger than the ideal compensation value. This could reflect premeditated commitment toward one of the two lateral targets, such that the hand attempted to launch at a substantial angle to the channel field. Critically, as these outliers could bias the results, we excluded all trials (∼10%) for which the normalized hand force exceeded the 95% confidence interval of the normalized hand-force distribution, which was computed across the single-target conditions of the training and test phases. This allowed us to obtain comparable distributions of compensatory hand-forces for the single and double-target conditions.

### Statistical analysis

To test the time-course of adaptation to the curl force-fields, we ran a two-way rmANOVA with experiment phase (3 levels: early-training, late-training, test) and target location (2 levels: central-target, lateral-target) as within-participant factors. To evaluate whether the hand-force was significantly different across the single-target and double-target conditions, we ran a one-way repeated measures ANOVA with target condition (3 levels: central-target, double-target, lateral-target) as a within-participant factor.

Separate one-sample t-tests (normal-distribution assumption verified by the Shapiro– Wilk test) were conducted to test whether the reaching dynamics in the double-target conditions were consistent with those for reaching the central (>0 adaptation values) or lateral targets (<0 adaptation values; see previous section for details).

We also sought to identify when different reaching dynamics could first be discriminated between task conditions during reaches. To this aim, we ran time series receiver operator characteristic (ROC) analysis on subject-wise averages of the hand-force traces that were time-locked to the target fill event. This allowed us to evaluate the time-based contrasts in reaching dynamics across conditions, relative to when the final target was made explicit. Separate ROC analyses were conducted for central versus double, lateral versus double, and central versus lateral target contrasts. For each pairwise comparison, we calculated the area under the ROC curves (AUC) whose values range from 0 to 1 (0.5 = chance discrimination; 1= perfect discrimination; 0 = incorrect discrimination). We set the threshold for discrimination at 0.7 and defined the “candidate” discrimination time as the time-point at which the AUC first exceeded and then remained above the threshold for at least 50ms. Finally, we ran a two-pieces “DogLeg” linear regression analysis around the candidate discrimination time to index the earliest deviation-from-chance of the time series ROC curve (Carroll et al., 2019; Contemori et al., 2021a, 2021b, 2022). The ROC analyses were conducted via a custom MATLAB script (version R2018b; The MathWorks, Inc., Natick, MA).

To test whether the target-goal uncertainty in double-target conditions affected motor decision-making processes, we ran a two-way rmANOVA with target condition (2 levels: single-target, double-target) and experiment phase (2 levels: baseline, test) as within-participant factors on both the RT and half-movement time results.

We also wanted to test whether RT and half-movement time were associated with the reaching direction in the double-target conditions. To this aim, we ran linear mixed-effects model (LMEM) analyses with the initial reach direction as the covariate and either the RT, or the half-movement time, as the dependent variable across random factors defined by subjects (24 levels). Note that this analysis was conducted solely on the no-force double-target trials of the baseline phase (i.e. when reach direction was not constrained by a force channel; Figure 1F).

The ANOVA, LMEM and t-test analyses were conducted in Jamovi (Jamovi project, v. 2.2.5). For the ANOVA analyses, we applied the Greenhouse-Geisser correction when the Mauchly’s test of sphericity was statistically significant. We also estimated the effect size by computing the Partial eta squared (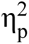). When the ANOVA revealed a significant main effect, we ran Bonferroni tests for post-hoc comparisons. For all tests, the statistical significance was designated at *p*<0.05.

## RESULTS

In this work, we revisited the question of how multiple options for action are represented within the neuromotor system. Specifically, we replicated the experimental protocol in Alhussein and Smith (2021) to induce target-dependent adaptation to curl force-fields but forced the participants to react in less than 500ms from the target presentation. We then tested which of the newly learnt reaching dynamics was adapted in circumstances requiring rapid decision amongst multiple potential targets for reaching.

### Intermediate-reaching behaviour under time pressure

Figure 2 shows the hand reaching trajectories of an exemplar participant. Unsurprisingly, almost straight target-directed trajectories were produced in the no-force-field trials (Figure 2A), while the reaching trajectory was perturbed toward the robot-force direction in the curl force-field conditions (Figure 2C). In both circumstances, trial-by-trial variability in reaching trajectory was removed when the hand movement was confined within the virtual mechanical channel (Figures 2B and D).

**Figure 2:**
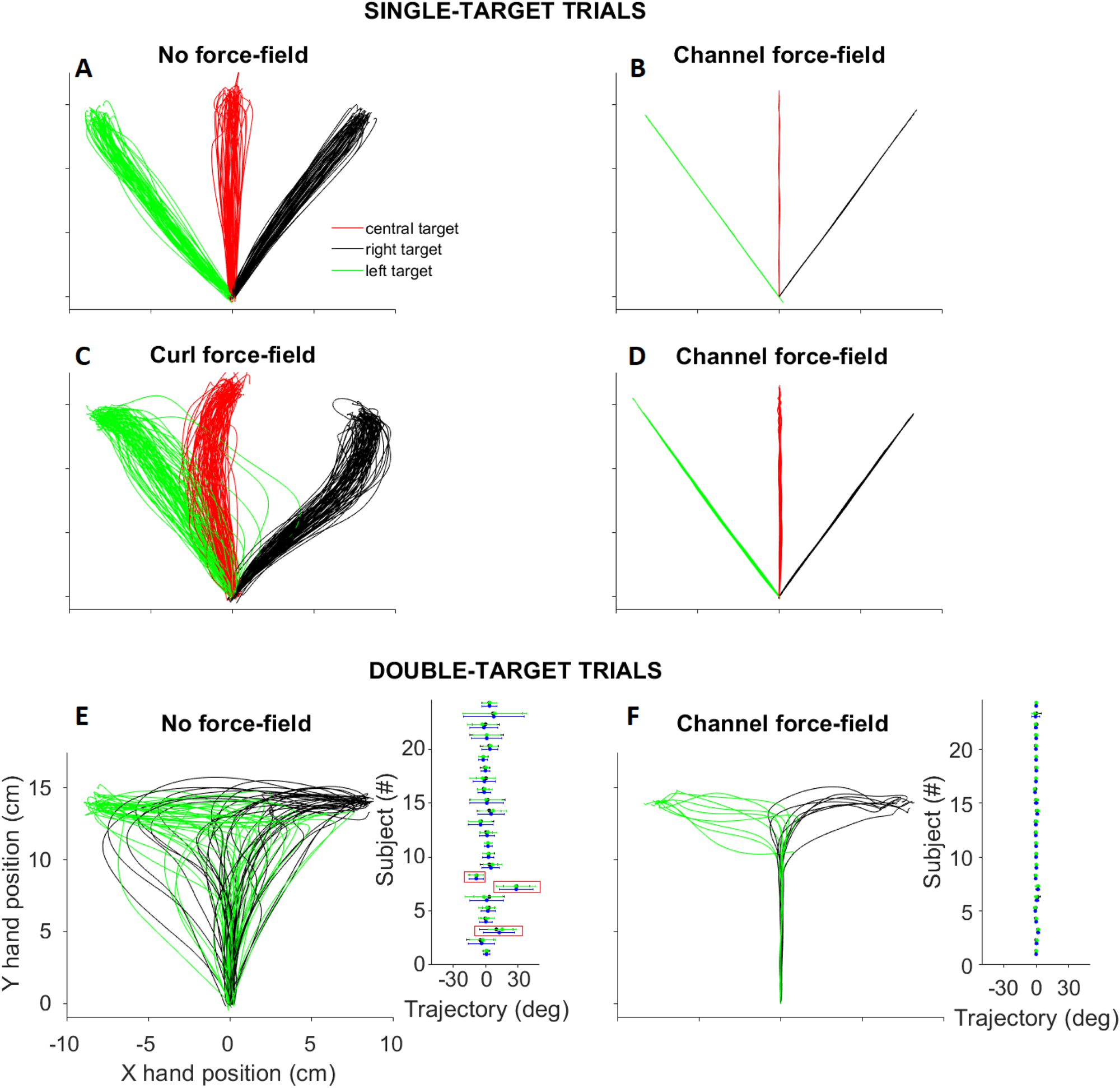
Reaching trajectories of an exemplar participant. A and B, single-target trials with no force-field and channel force-field of the baseline phase. C and D, single-target trials with curl force-field and channel force-field of the training phase. In this example, CCW an CW force-fields were applied for the central and lateral targets, respectively. E, double-target trials with no force-field. E and F, double-target trials with no force-field and channel force-field of the baseline phase. The inset plots show the average (dot) and standard deviation (error bar) of the initial reach direction in the double-target conditions for each of the 24 subjects (colour coding: green for the right target; black for the left target; blue for the average across right and left targets). The red frames indicate the subjects (N=3) who had target-dependent biases for reaching as they systematically reached toward one of the two potential targets. Each trajectory trace in panels A-F represents a single trial completed by participant #17 in panels E and F.

In double-target conditions, movements were initiated prior to knowing the final target to reach. Given this uncertainty, participants sometimes opted to reach one of the two targets on the no-force double-target trials of the baseline phase although mismatching the final target required to change the reaching direction (see the outermost traces in Figure 2E). In most of the trials, however, the reach was initially directed toward an intermediate location between the possible targets (Figure 2E). Thus, the initial reach-direction angle was distributed around 0° (see inset plot of Figure 2E). Also, the reaching redirection toward the final target is reflected by the biphasic hand-speed traces in the double-target conditions (Figures 3A-B).

**Figure 3:**
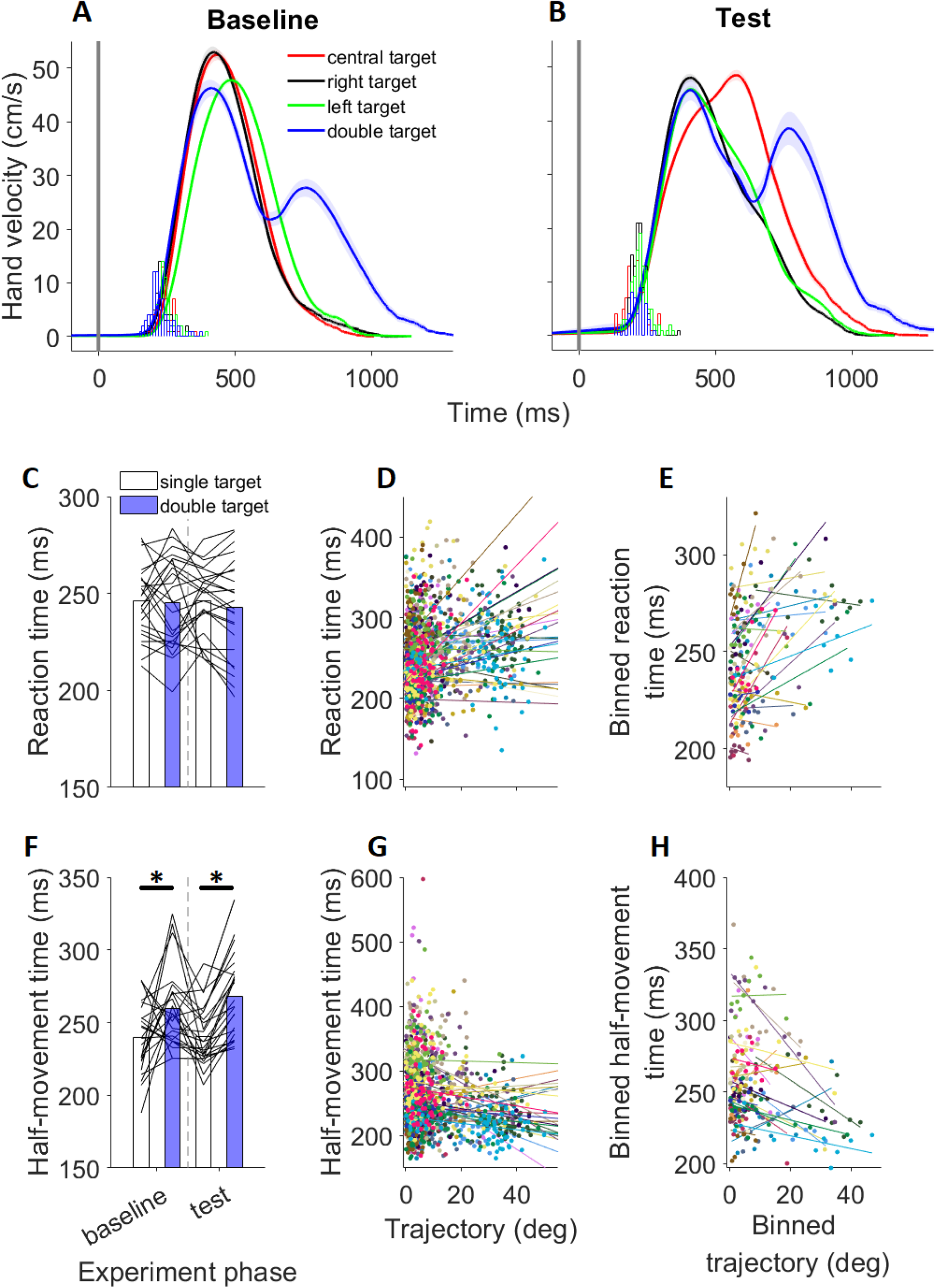
A-B, Hand-velocity traces for the exemplar participant featured in figure 2 across the different task conditions in (A) the baseline and (B) test phases. The data are shown as means (solid lines) and standard errors (shaded areas around the mean lines) across trials. The histograms represent the condition-based distribution of the RT. The data are plotted relative to the target presentation time (grey vertical line at 0ms). C, Dependency of the RT on the target condition (single-target VS double-target) and experiment phase (baseline VS test). Each line represents one subject, and the bars represent the average across subjects. D, Subject-specific relationship between the initial reach direction (0 deg = midline reach direction) and the RT for the and experiment phase (2 levels: baseline, test) as within-participant factors on both the RT and half-movement time results. Each subject is represented via a different colour, each data point represents a single trial, and each line represents the linear model fitted to the data. E, Subject-specific relationship between the initial reach direction and the RT for the binned data samples (9 bins) from the smallest to the largest angle of the initial reach trajectory (same format as panel D). F, Dependency of the half-movement time on the target condition and experiment phase (same format as panel C). The horizontal black lines and asterisks represents the statistically significant (p<0.05) contrasts. D and H, Subject-specific relationship between the initial reach direction and the half-movement time for (D) the original and (H) binned data samples (same format as panels D and E).

Similar results were found among 21 of the 24 subjects, whereas three participants exhibited target-dependent biases such that they tended to systematically reach toward one of the two potential targets in the double-target conditions (right-target for participants #3 and #7; left-target for participant #8; inset plot in Figure 2E). Note that heterogeneity of intermediate reaching behaviour is consistent with previous work showing that participants flexibly adapt or abandon it depending on the surrounding context (Wong et al., 2017, 2022). Most of our participants, however, (i.e. 87.5%) adopted the intermediate-reaching behaviour even under the imposed time pressure on motor decision-making processes.

### Target-goal uncertainty did not affect the reaction time

For the exemplar subject in Figure 2, the distribution of RTs appears similar across the single-target and double-target conditions (Figures 3A-B), suggesting that the time to start moving was independent of advance knowledge about the final target to reach for this person. We used a bootstrap approach to exclude significant differences in RT between the different target locations across the three experiment phases for any of the participants. Having excluded target-dependent RT differences, and then collapsed RTs across the three single-targets and for a contrast between the single-target and double-target conditions for the baseline and test phases. We found similar RT across the double-target and single-target conditions (target condition main-effect: *F_1,23_*=0.7, *p*=0.42, 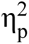=0.028), and for both the baseline and test phases (experiment phase main-effect: *F_1,23_*=0.1, *p*=0.75, 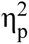=0.04; target*phase interaction: *F_1,23_*=0.55, *p*=0.46, 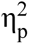=0.024). This suggests that movement onset time was independent of prior knowledge about the final goal for action.

### Earlier initiation of intermediate than direct reaches in double target trials

A LMEM analysis returned a positive and significant linear relationship between RT and reaching direction, both for all individual trials (*F_1,23_*=19.6, *p*<0.001, estimated slope=0.83, R^2^=0.31; Figure 3D) and for nine data points obtained by binning the RT and reaching-direction data from the smallest to the largest reach-trajectory angle (*F_1,23_*=19.9, *p*<0.001, estimated slope=0.93, R^2^=0.8; Figure 3E). Further, we excluded that any association with reductions in RT over time. In fact, while a negative and significant relationship existed between the trial number and the RT (*F_1,23_*=4.4, *p*=0.046, estimated slope=-0.14, R^2^=0.29), no significant relationship was found between the trial number and the initial reaching direction (*F_1,23_*=0.24, *p*=0.63, estimated slope=-0.007, R^2^=0.43). Overall, these results indicate that intermediate reaches were initiated systematically earlier than those that were more directed toward either of the two alternative targets.

Our data also show that the reaching velocity was lower for the double-target than single-target conditions. The rmANOVA analysis, indeed, returned a significant target-condition effect on the half-movement time (*F_1,23_*=28.3, *p*<0.001, 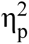=0.55), while neither a significant phase-dependent modulation (*F_1,23_*=1.6, *p*=0.22, 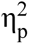=0.07), nor a target*phase interaction (*F_1,23_*=0.86, *p*=0.36, 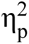=0.04) were found. The post-hoc tests showed that the half-movement time was significantly longer for the double-target than the single target conditions for both the baseline and test phases (Figure 3F). This indicates that the reaching action evolved at a slower velocity for intermediate reaches than for target-directed reaches, a result that is also corroborated by the LMEM analysis. In fact, we found a negative and significant linear relationship between the initial reach direction and half-movement time for both the original (*F_1,23_*=9.64, *p*=0.005, estimated slope=-0.66, R^2^=0.31) and binned (*F_1,23_*=12, *p*=0.003, estimated slope=-0.74, R^2^=0.74) datasets (Figures 3G and H).

### Suboptimal adaptation under time pressure

Figure 4A shows the group-averaged (N=17) adaptation to the target-dependent FFs along the different experiment phases. Note that seven subjects were excluded from this analysis as they did not adapt properly to the curl force-fields. Specifically, these subjects adopted the newly learnt reaching dynamics for the lateral target across all the target locations, while three of them also tended to systematically reach toward one of the two lateral targets in the baseline double-target conditions (Figure 2E). Critically, this happened despite the replication of a testing protocol that was proven effective to induce appropriate FF-adaptations among all participants (Alhussein and Smith, 2021). It is worth noting, however, that Alhussein and Smith (2021) cued the movement onset 1000ms after target presentation and then allowed an additional 425ms to start moving, whereas we allowed only 500ms to react (see Materials and Methods).

**Figure 4:**
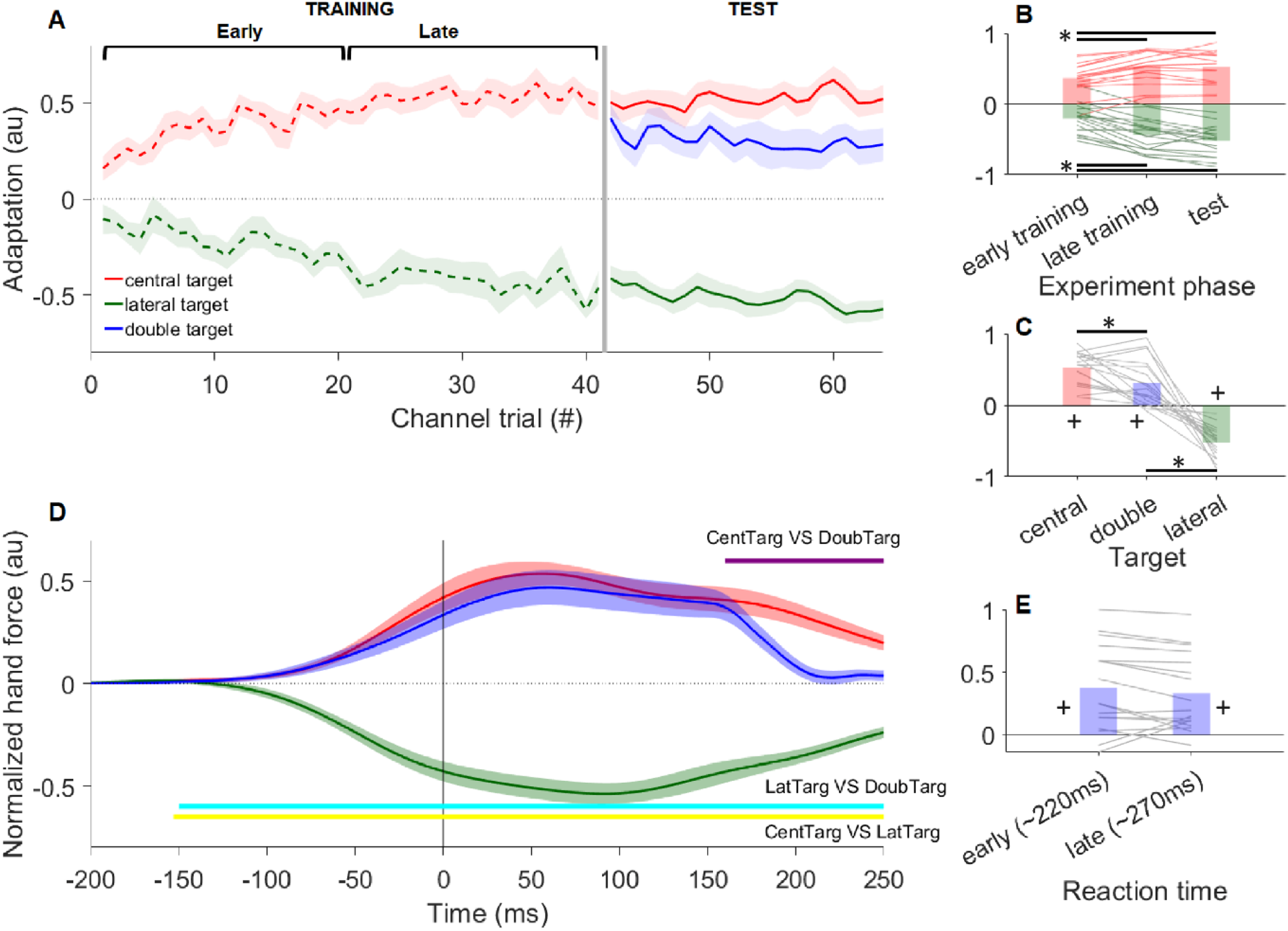
A, Time-course of adaptation to the target-dependent FFs. B, Dependency of the adaptation to the central-target (red) and lateral-targets (green) FFs among the early-training, late-training, and test phases of the experiment. The horizontal black lines and asterisks represent statistically significant (p<0.01) differences between the experiment phases. C, Dependency of adaptation magnitude at the test phase on the central-target, lateral-target, and double-target (blue) conditions. The horizontal black lines and asterisks represents statistically significant (p<0.05) differences between the target conditions; the ‘+’ symbols represent the target conditions for which the amount of adaptation was significantly different (p<0.05) from zero. D, Temporal evolution of the hand-force for the different target conditions relative to the target fill event (vertical black line at 0ms). The horizontal lines show the results of pairwise (central_VS_double; central_VS_lateral; lateral_VS_double) ROC analyses and indicate the time points at which the target conditions were first discriminable solely from the hand-force trace. E, Adaptation magnitude in the double-target conditions across fast and slow RT subsets. In panels A and D, the data are shown as means (solid lines) and standard errors (shaded areas around the mean lines) across subjects (N=17). In Panels B, C and E, each line represents one subject, and the bars represent the average across subjects.

For the seventeen subjects who adapted properly, the amount of adaptation increased with prolonged exposure to the opposite target-dependent fore-fields (target-location effect: *F_1,16_*=3.16, *p*=0.09, 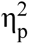=0.17; experiment-phase effect: *F_2,32_*=58, *p*<0.001, 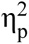=0.78). On average, around half of the ideal level of compensating force was applied by the end of the training phase and remained largely preserved during the test phase (Figure 4A-B). We also found a statistically significant interaction effect between target-location and experiment-phase (*F_2,32_*=5.7, *p*=0.013, 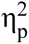=0.6). The force-field adaptation, indeed, increased between the late-training and test phases for the lateral-target (−0.43 → −0.52), whereas it remained similar for the central target (∼0.53; Figure 4B). This could be due to the larger prevalence of lateral-target than central-target trials presented during the test phase (see Materials and Methods and figure 1F for details).

Overall, these results indicate that most of the participants were able to appropriately adapt to the opposite perturbing force-fields despite the imposed time pressure on motor decision-making processes.

### The reaching dynamics of intermediate reaches compensate the force-field for the central rather than the lateral targets

The main finding of the current work is that the reaching dynamics in the double-target conditions were compatible with those learnt to compensate the curl force-field applied during central-target reaches (Figure 4A). The one-sample t-tests, indeed, showed that the adaptation was significantly larger than zero for the central-target (*p*<0.001) and double-target (*p*=0.001) conditions, whereas it was significantly lower (*p*<0.001) than zero for the lateral targets (Figure 4C). Notably, these results held true also for trials with short reaction times as indicated by a median split analysis on the RT values (Figure 4E; see table 1 for subject-wise RT split details).

**Table 1:**
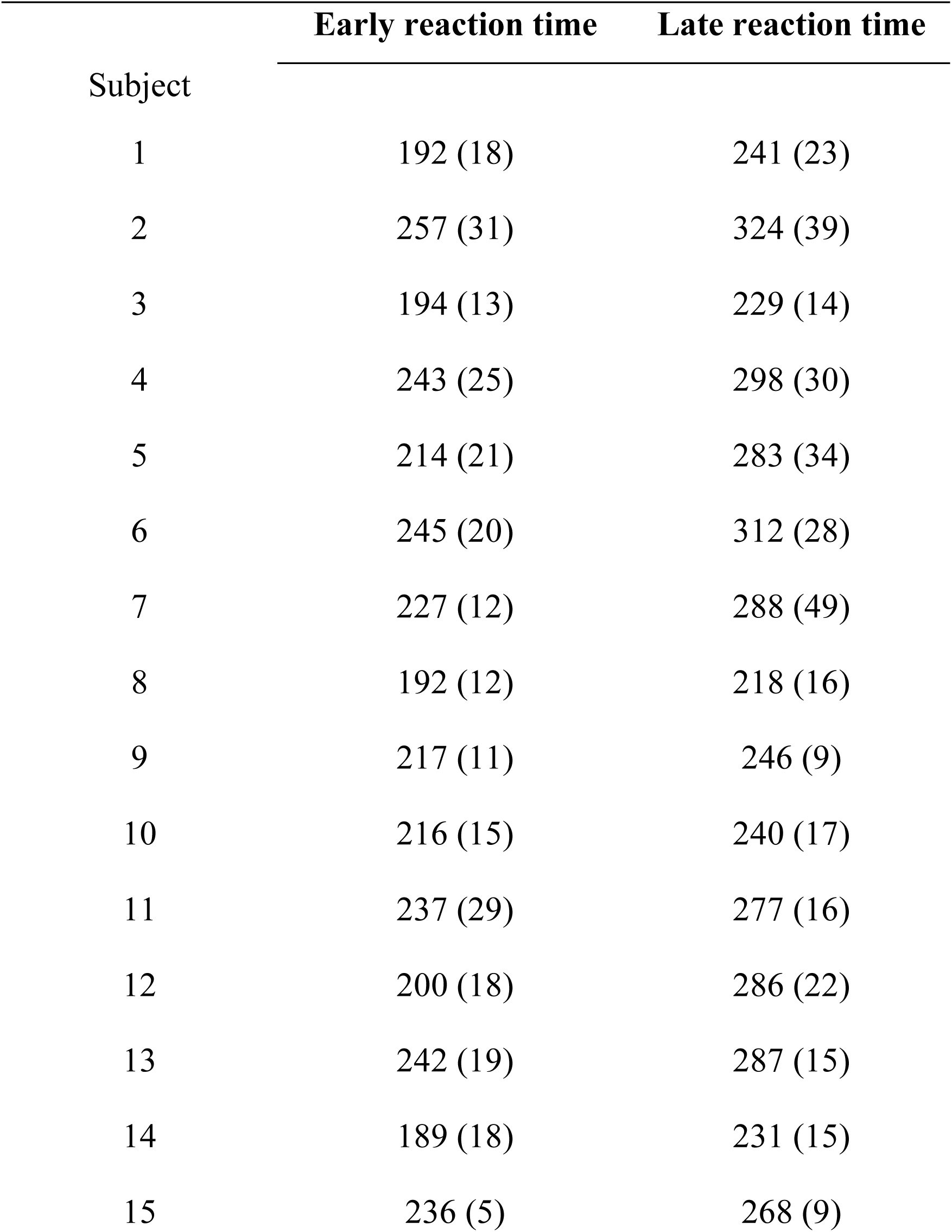
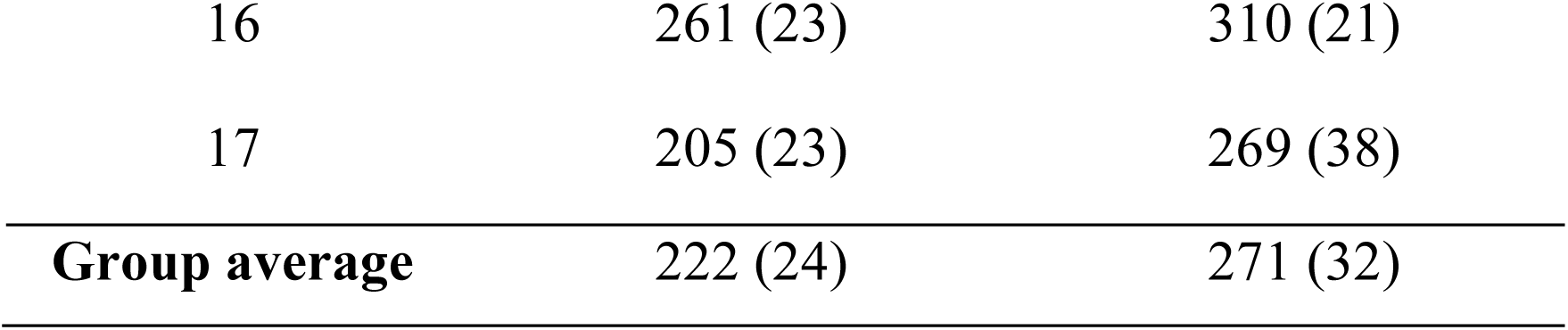
Average (standard deviation) reaction time computed across the trials for which the reaction time was earlier and later than the median values.

| Subject | Early reaction time | Late reaction time |
| --- | --- | --- |
| 1 | 192 (18) | 241 (23) |
| 2 | 257 (31) | 324 (39) |
| 3 | 194 (13) | 229 (14) |
| 4 | 243 (25) | 298 (30) |
| 5 | 214 (21) | 283 (34) |
| 6 | 245 (20) | 312 (28) |
| 7 | 227 (12) | 288 (49) |
| 8 | 192 (12) | 218 (16) |
| 9 | 217 (11) | 246 (9) |
| 10 | 216 (15) | 240 (17) |
| 11 | 237 (29) | 277 (16) |
| 12 | 200 (18) | 286 (22) |
| 13 | 242 (19) | 287 (15) |
| 14 | 189 (18) | 231 (15) |
| 15 | 236 (5) | 268 (9) |
| 16 | 261 (23) | 310 (21) |
| 17 | 205 (23) | 269 (38) |
| **Group average** | 222 (24) | 271 (32) |

| Subject | Early reaction time | Late reaction time |
| --- | --- | --- |
| 1 | 192 (18) | 241 (23) |
| 2 | 257 (31) | 324 (39) |
| 3 | 194 (13) | 229 (14) |
| 4 | 243 (25) | 298 (30) |
| 5 | 214 (21) | 283 (34) |
| 6 | 245 (20) | 312 (28) |
| 7 | 227 (12) | 288 (49) |
| 8 | 192 (12) | 218 (16) |
| 9 | 217 (11) | 246 (9) |
| 10 | 216 (15) | 240 (17) |
| 11 | 237 (29) | 277 (16) |
| 12 | 200 (18) | 286 (22) |
| 13 | 242 (19) | 287 (15) |
| 14 | 189 (18) | 231 (15) |
| 15 | 236 (5) | 268 (9) |
| 16 | 261 (23) | 310 (21) |
| 17 | 205 (23) | 269 (38) |
| **Group average** | 222 (24) | 271 (32) |

The adaptation was significantly larger than zero (*p*<0.001) for both the fast and slow subsets (Figure 4E), with no significant between-subset difference (paired sample t-test: *p*=0.09). The direction of the non-significant trend indicated that, if anything, the shorter reaction times tended to result in stronger adaptation magnitudes appropriate for the central target. Thus, even at the shortest reaction times, our results align with those from Alhussein and Smith (2021).

The conclusion that the reaching dynamics of intermediate reaches reflected only those learnt for reaching toward the central target is corroborated by the results of the ROC analysis (Figure 4D). Specifically, we found that the reaching dynamics for the central-target and double-target conditions were not significantly discriminable up to 150ms after the target fill event, a time consistent with the removal of the mechanical channel in the double-target conditions (see Materials and Methods and Figure 1D for details). By contrast, the dynamics for reaching either of the lateral targets were discriminable from both the central-target and double-target dynamics well before (∼150ms) the final target was presented (Figure 4D).

As shown in figure 4C, the adaptation was significantly lower in the double-target than single target conditions (*F_2,32_*=7.3, *p*=0.004, 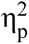=0.31; central_VS_double *p*=0.033, lateral_VS_double *p*=0.012). These findings, however, were again consistent with the results of Alhussein and Smith (2021), and may reflect variability in intended initial reaching direction in the double-target condition (Figure 2E).

## DISCUSSION

In this work, we exposed human subjects to target-dependent curl force-fields such that distinct reaching dynamics were learned for central and lateral targets. We showed that people applied the newly learnt reaching dynamics for the central target when the two lateral targets appeared simultaneously in the “go before you know” paradigm. Importantly, these results were obtained by imposing time pressure on decision-making processes and, thereby, limiting the opportunity for cognitive deliberation among different action alternatives. The results are consistent with the interpretation that reach dynamics were specified downstream of decisions about where to move in our task. The data are relevant to ongoing debate about whether the brain decides on a unique action from multiple target alternatives before producing a single motor plan, or partially prepares multiple motor actions in parallel.

### Dynamics of reaching are not averaged even under time pressure

Our result show that the dynamic features of reaching actions were consistent with those required for a previously visited central target rather than those needed for a combination of presented (lateral) targets. These findings mirror those of Alhussein and Smith (2021), who used the same curl field manipulations as in the current study to dissociate the required reach dynamics for alternative targets, and Nashed et al. (2017), who used simulated spring dynamics requiring grip force modulations for different target directions. However, Alhussein and Smith allowed participants more than 1 second to view potential targets before reaching, and although Nashed et al. prompted movement initiation at the same time as target presentation, there were no reaction time constraints, and reactions times were not reported. Thus, participants had opportunity to deliberate about the best action to take under uncertainty in both of these studies. Instead, reach direction in the current study is more likely to reflect low-level sensorimotor processes to integrate potential target information under goal uncertainty. Taken together, it appears that participants do not “average” dynamic features of reaching actions between targets that are presented as potential alternatives in “go before you know” tasks, even when there is limited time for deliberation about how to respond to the target uncertainty.

It is important to consider how effective our time limit methods were. Our participants were forced to initiate movement within 500ms of target presentation and, in fact, produced reactions with a substantial safety margin of this hard constraint (i.e. grand mean RT calculated post hoc < 250ms). This safety margin presumably exists because the online movement initiation threshold we used (i.e. the time when the cursor exited the start circle) lags the actual movement initiation time as detected by a more sensitive method based on hand speed measurements, and because participants were averse to the error message and/or need to repeat trials that were initiated too late. In any case, our observed reaction times were short compared to typical multi-target reaching tasks (e.g. Li et al., 1995; Gribble et al., 2002; Fernandez-Ruiz, 2010) Further, although people can voluntarily re-aim their reach directions away from a target in less than 250ms (down to ∼150ms; Leow et al. 2017), such re-aiming comes at a considerable accuracy cost and requires focused concentration and effort. Moreover, the extent to which participants corrected for the imposed force field on single target trials was considerably lower in the current study than in an equivalent task without time pressure (i.e. ∼50% here VS ∼100% in Alhussein and Smith, 2021), implying that the reaction time constraints in our study dramatically affected motor learning behaviour. Crucially, the reaching dynamics were not significantly different between the shortest and longest reaction time trials for our double target condition, which suggests that a tighter reaction time constraint would have been unlikely to alter the core findings of the study. We conclude that our reaction time constraint provided an effective pressure for participants to respond as quickly as possible and limited the opportunity for “cognitive” deliberation about the best action to produce in the period between target presentation and movement initiation.

### Competition between multiple motor plans or a single plan optimised for task performance?

Although our data support the conclusion that multiple options for action are represented via a single motor plan (Nashed et al., 2017; Alhussein and Smith, 2021), it is important to consider whether this proposal is generally applicable beyond the task context of the relevant experiments. For example, Enachescu et al. (2021) presented a normative model for action decisions that illustrates how behaviour ranging from direct reaches to one of two potential targets, to intermediate reaches between targets, can be produced by a control architecture involving competition and averaging between movement-control policies. The key aspect of the model that allows for a broad range of behaviour depending on context is the incorporation of a composite “desirability” integrator that weights potential actions according to their values and costs. This aspect predicts direct-to-one-target reach behaviour in “go before you know” tasks that was previously interpreted as providing evidence for a single motor plan with performance optimisation (e.g. Haith et al., 2015a; Wong and Haith, 2017; Wong et al., 2022). Specifically, the model predicts direct reaching to one specific target when target options are spatially distant, because corrective movement costs are much higher in this case, and also predicts direct to target reaching when barriers prevent intermediate movements, because the value of intermediate movements is low. Thus, a failure to observe motor averaging in a specific context does not provide definitive evidence that the mechanism for action selection under uncertainty does not involve competition among “movement plans” for alternative options.

The capacity for the normative model of Enachescu et al. (2021) to predict the current and previous (Nashed et al. 2017; Alhussein and Smith 2021) results depends on the assumption that internal representations of actions to potential targets that are not physically displayed can compete for selection with actions to the presented targets. Under this assumption, intermediate movements between two presented targets that apply reach dynamics for an absent central target could occur because a “reach plan” for the central target reaches threshold before either of the two lateral targets, and because the value of the lateral target reach plans is low because they are incompatible with the central reach plan (i.e. leading to winner takes all behaviour for reach dynamics of the central target). In support of this idea is our observation of shorter reaction times for intermediate movements than for movements directed to either lateral target. This suggests that there is a bias toward preparation for the central target and can be explained by competition among reach plans assuming that competition is resolved fastest when the internal representation of the (most valuable) central target is strong. The representation of a “virtual” central target makes sense given the designs of the current study and those of Nashed et al. (2017) and Alhussein and Smith (2021), because participants received extensive experience reaching to central targets. Moreover, during the test phases in which reach dynamics were assessed, any of the three individual targets (left, right, central) could be displayed on a given trial.

Making distinction between a competition among motor plans to reach physical and virtual targets and the “performance optimisation” hypothesis may seem a trivial exercise in semantics, given that the outcomes of both hypotheses in this case resembles a motor plan that is specifically optimised for the task, rather than a weighted average of plans for the presented targets. However, there is a crucial mechanistic principle at stake, namely: at what stage in the sensorimotor transformation process does the brain (or more generally, the CNS) decide among various options for action? The performance optimisation hypothesis assumes a higher order decision occurs before an action is planned, whereas in the motor encoding model, decision making occurs through competition between alternative actions at all levels of sensorimotor representation – including motor planning. If decision making is to effectively consider the costs of various action alternatives, then an internal representation of alternative costs should contribute to the decision process. This could be efficiently achieved by recurrent connections between areas that specify the movement details (e.g. Cisek & Kalaska, 2010; Enachescu et al., 2021; Hunt & Hayden, 2017; Steinmetz et al., 2019; Rao, 2024), but could also occur by independent representations of motor costs associated with movements in higher order decision making brain areas (e.g. Haith et al., 2015a; Wong and Haith, 2017; Alhussein and Smith, 2021; Wong et al., 2022). These alternative conceptions speak to the neural implementation of reach and choice behaviour under uncertainty.

### Neural evidence for planning of more than one action at a time?

There is an extensive literature comprised of experiments that used single electrode intracortical recordings that was interpreted as evidence that information related to multiple potential actions is encoded in sensorimotor and decision-making brain regions (see Cisek & Kalaska, 2010; Hunt & Hayden, 2017; Gallivan et al. 2018 for review). However, Dekleva et al. (2018) recently showed that reconstructions of brain representations from multiple trials and sessions can provide misleading support for dual representations in situations in which there are alternating discrete representations from trial to trial. They considered activity in dorsal premotor cortex and primary motor cortex of monkeys engaged in a “go before you know” task and found that multi-unit dynamics associated with movement preparation always reflected only one of the two alternative targets on any given trial. This was despite replicating apparent dual encoding of two actions via ensemble representations based on trial averages. They concluded that motor cortical areas represent only a single motor plan at a time, in agreement with the performance optimisation hypothesis. However, the task in this study likely favoured single action planning, as the two alternatives were always 180 degrees apart, and therefore incompatible for averaging. Also, on a subset of trials, the monkeys were cued to make a free choice about which target to reach immediately after the trial period used for analysis of preparation dynamics, which might also have encouraged them to choose one of the two options prior to the go signal. Thus, although there was no evidence of competing motor plans in this particular context, it remains possible that action encoding mechanisms occur in motor cortical areas under different task conditions. Indeed Meirhaeghe et al. (2023) found dynamical states of preparatory neural activity that were intermediate between those associated with two alternative grasping movements in premotor cortex. Also, Siegel et al (2015) reported choice-related activity recorded simultaneously in multiple sensorimotor cortical areas in monkeys, which was consistent with a distributed decision-making mechanism. Finally, Steinmetz et al. (2019) recorded simultaneously from 30,000 neurons in 42 brain regions of mice performing a visual discrimination task and found choice-related signals in both frontal cortex and the midbrain that evolved simultaneously, suggesting that decision making occurs by competition of action representations via recurrent connections between multiple brain areas. This evidence suggests that competition between partially prepared reach plans remains a viable mechanism for action selection under goal uncertainty in reaching - but what about the specific case of reach dynamics specification?

### Where are the movement dynamics computed?

Perich and colleagues (2018) showed that adaptation to perturbing force fields led to modification of the output-null (or preparatory) dimension of dynamical states constructed from neural activity in the dorsal premotor area. They concluded that new movement dynamics are learnt by changing the instructions to a downstream area that computes movement dynamics and drives the muscle action. Although it is generally assumed that primary motor cortex is the key downstream region for reach dynamics computation, an alternative perspective based on a hierarchical conception of sensorimotor control which emphasises that multiple downstream areas including the brainstem and spinal cord might contribute to computations of limb dynamics (e.g. Loeb, 2012; Contemori et al., 2023; Rao, 2024). In this case, preparatory cortical activity would set the states of subcortical circuits to transform descending control signals into task-appropriate motoneuron drives. For example, simulations of spinal reflex circuits show that proprioceptors and known interneuronal connections can entirely compensate for imposed force fields during reaching on the basis of alterations in pre-movement gain settings (Raphael et al, 2010; Tsianos et al, 2014). Importantly, the simulations also showed that appropriate muscle activations for novel actions can be achieved by interpolation of subcortical state settings for similar targets that were previously encountered, implying that motor averaging of reach dynamics is feasible at the spinal level.

Behavioural evidence for the contribution of multiple downstream areas to movement dynamics specification comes from observations of appropriately goal-directed “express” muscle activations at latencies approaching the minimal sensorimotor conduction delay times (<100ms), suggesting involvement of the superior colliculus, reticular formation (Pruszynski et al, 2010; Gu et al, 2016; Contemori et al., 2022, 2023), and spinal cord (Weiler et al., 2019, 2021). Indeed, Selen et al. (2023) showed that “express” muscle activations were tuned to drive the limb to an intermediate direction between alternative targets in a free choice reaching task. This suggests competition between alternative actions can occur at low-level, likely subcortical, stages of sensorimotor transformation and is consistent with the notion of distributed motor decision making. By contrast, the current data suggest that action selection occurs upstream of the circuits that specify reach dynamics. This might have occurred because the dynamics of the two lateral targets were incompatible with those of the central target. In such a scenario, gain setting for subcortical circuit states might reflect one or the other target because of reciprocal inhibition. If this is the case, reaction time data from the current study would suggest that states appropriate for the different dynamics associated with different targets can be switched rapidly (∼20ms).

## Conclusion

The key finding of this study was that reach dynamics were not averaged for alternative targets presented as options in a “go before you know” task, even when participants were placed under time pressure to respond as fast as possible. Although this indicates that reach dynamics were specified downstream of decisions about where to move in this particular task, it remains an open question whether such behaviour arises from the generation of a single motor plan that is optimised for task performance based on higher order decision signals, or from competition between alternative motor plans that was resolved upstream of reach dynamics specification.

## Acknowledgements

The work was supported by Australian Research Council grant DP230102179. We thank Reza Shadmehr for helpful suggestions.

## REFERENCES

Alhussein L, Smith MA. Motor planning under uncertainty. eLife 10:e67019, 2021.

Ames KC, Ryu SI, Shenoy KV. 2019. Simultaneous motor preparation and execution in a last-moment reach correction task. Nat. Commun. 10(1):2718.

Ames, C. K., Ryu, S. I. & Shenoy, K. V. Neural dynamics of reaching following incorrect or absent motor preparation. Neuron 81, 438–451 (2014).

Baluch F, Itti L. Mechanisms of top-down attention. Trends Neurosci 34: 210–224, 2011.

Billen LS, Corneil BD, Weerdesteyn V (2023). Evidence for an intricate relationship between express visuomotor responses, postural control and rapid step initiation in the lower limbs. Neuroscience 531:60–74.

Brenner E, de la Malla C, Smeets JBJ (2022) Tapping on a target: dealing with uncertainty about its position and motion. Exp Brain Res 241:81–104.

Carroll TJ, McNamee D, Ingram JN, Wolpert DM. Rapid visuomotor responses reflect value-based decisions. J Neurosci 39: 3906–3920, 2019

Chapman CS, Gallivan JP, Wood DK, Milne JL, Culham JC, Goodale MA. 2010. Reaching for the unknown: multiple target encoding and real-time decision-making in a rapid reach task. Cognition 116:168–176.

Cisek P, Kalaska JF. 2005. Neural correlates of reaching decisions in dorsal premotor cortex: specification of multiple direction choices and final selection of action. Neuron 45:801–814.

Cisek P. 2007. Cortical mechanisms of action selection: the affordance competition hypothesis. Philosophical Transactions of the Royal Society B: Biological Sciences 362:1585–1599.

Connolly JD, Goodale MA, DeSouzaJF, Menon RS, Vilis T. 2000. A comparison of frontoparietal fMRI activation during anti-saccades and anti-pointing. J Neurophysiol. 84:1645–1655.

Contemori S, Loeb GE, Corneil BD, Wallis G, Carroll TJ (2021a) The influence of temporal predictability on express visuomotor responses. J Neurophysiol 125:731–747.

Contemori S, Loeb GE, Corneil BD, Wallis G, Carroll TJ (2021b) Trial-by-trial modulation of express visuomotor responses induced by symbolic or barely detectable cues. J Neurophysiol126(5):1507–1523.

Contemori S, Loeb GE, Corneil BD, Wallis G, Carroll TJ (2022) Symbolic cues enhance express visuomotor responses in human arm muscles at the motor planning rather than the visuospatial processing stage. J Neurophysiol 128(3):494–510.

Contemori S, Loeb GE, Corneil BD, Wallis G, Carroll TJ (2023). Express visuomotor responses reflect knowledge of both target locations and contextual rules during reaches of different amplitudes. J Neurosci JN-RM-2069–22, epub ahead of print. DOI: 10.1523/JNEUROSCI.2069-22.2023.

Crevecoeur F, Thonnard JL, Lefèvre P (2020a) A very fast time scale of human motor adaptation: within movement adjustments of internal representations during reaching. eNeuro 7:ENEURO.0149-19.2019.

Crevecoeur, F., Mathew, J., Bastin, M., and Lefèvre, P. (2020b). Feedback adaptation to unpredictable force fields in 250 ms. eNeuro 7:ENEURO.400-19.2020.

Cross KP, Cluff T, Takei T, Scott SH (2019) Visual feedback processing of the limb involves two distinct phases. J Neurosci 39:6751–6765.

Day BL, Brown P (2001) Evidence for subcortical involvement in the visual control of human reaching. Brain 124:1832–1840.

Day BL, Lyon IN (2000) Voluntary modification of automatic arm movements evoked by motion of a visual target. Exp Brain Res 130:159–168.

de Xivry, J.-J., Legrain, V. & Lefèvre, P. Overlap of movement planning and movement execution reduces reaction time. J. Neurophysiol. 117, 117–122 (2016).

Dekleva BM, Kording KP, Miller LE. 2018. Single reach plans in dorsal premotor cortex during a two-target task. Nature Communications 9:3556.

Enachescu V, Schrater P, Schaal S, Christopoulos V (2021) Action planning and control under uncertainty emerge through a desirability-driven competition between parallel encoding motor plans.

Everling S, Fischer B. (1998). The antisaccade: A review of basic research and clinical findings. Neuropsychologia, 36, 885–899.

Fautrelle L, Prablanc C, Berret B, Ballay Y, Bonnetblanc F (2010) Pointing to double-step visual stimuli from a standing position: very short latency (express) corrections are observed in upper and lower limbs and may not require cortical involvement. Neuroscience 169:697–705.

Fernandez-Ruiz J, Wong W, Armstrong IT, Flanagan JR. Relation between reaction time and reach errors during visuomotor adaptation. Behav Brain Res 219: 8–14, 2011.

Forano M, Schween R, Taylor JA, Hegele M, Franklin DW. Direct and indirect cues can enable dual adaptation, but through different learning processes. J Neurophysiol 2021;126(5):1490–1506.

Gallivan JP, Barton KS, Chapman CS, Wolpert DM, Flanagan JR. 2015. Action plan co-optimization reveals the parallel encoding of competing reach movements. Nature Communications 6:7428. DOI: 10.1038/ncomms8428, PMID: 26130029

Gallivan JP, Chapman CS, Wolpert DM, Flanagan JR. 2018. Decision-making in sensorimotor control. Nat. Rev. Neurosci. 19(9):519–34.

Gallivan JP, Chapman CS, Wood DK, Milne JL, Ansari D, Culham JC, Goodale MA. 2011. One to four, and nothing more: nonconscious parallel individuation of objects during action planning. Psychological Science 22:803–811. DOI: 10.1177/0956797611408733, PMID: 21562312

Gallivan JP, Chapman CS. 2014. Three-dimensional reach trajectories as a probe of real-time decision-making between multiple competing targets. Frontiers in Neuroscience 8:215.

Gallivan JP, Logan L, Wolpert DM, Flanagan JR. 2016. Parallel specification of competing sensorimotor control policies for alternative action options. Nature Neuroscience 19:320–326. DOI: 10.1038/nn.4214, PMID: 26752159

Gallivan JP, Stewart BM, Baugh LA, Wolpert DM, Flanagan JR. 2017. Rapid automatic motor encoding of competing reach options. Cell Reports 18:1619–1626.

Gribble PL, Everling S, Ford K, and Mattar A. Hand-eye coordination forrapid pointing movements. Arm movement direction and distance are specified prior to saccade onset. Exp Brain Res 145: 372–382, 2002.

Gu C,Wood DK, Gribble PL, Corneil BD (2016) A trial-by-trial window into sensorimotor transformations in the human motor periphery. J Neurosci 36:8273–8282.

Haith AM, Huberdeau DM, Krakauer JW. 2015a. Hedging your bets: intermediate movements as optimal behavior in the context of an incomplete decision. PLOS Computational Biology 11:e1004171.

Haith AM, Huberdeau DM, Krakauer JW. The influence of movement preparation time on the expression of visuomotor learning and savings. J Neurosci 35: 5109–5117, 2015b.

Haith, A. M., Pakpoor, J. & Krakauer, J. W. Independence of movement preparation and movement initiation. J. Neurosci. 36, 3007–3015 (2016).

Howard IS, Ingram JN, Franklin DW, Wolpert DM. Gone in 0.6 seconds: The encoding of motor memories depends on recent sensorimotor states. J Neurosci 32:12756–12768, 2012.

Howard S, Ingram JN, Wolpert DM (2009) A modular planar robotic manipulandum with endpoint torque control. J Neurosci Methods 181(2):199–211.

Hunt, L.T., and Hayden, B.Y. (2017). A distributed, hierarchical and recurrent framework for reward-based choice. Nat. Rev. Neurosci. 18, 172–182.

Kaufman MT, Seely JS, Sussillo D, Ryu SI, Shenoy KV, Churchland MM. 2016. The largest response component in the motor cortex reflects movement timing but not movement type. eNeuro 3(4):ENEURO.0085–16.2016.

Kaufman, M.T., Churchland, M.M., Ryu, S.I., and Shenoy, K.V. (2014). Cortical activity in the null space: Permitting preparation without movement. Nat Neurosci. 17, 440–448.

Kozak RA, Kreyenmeier P, Gu C, Johnston K, Corneil BD (2019) Stimulus-locked responses on human upper limb muscles and corrective reaches are preferentially evoked by low spatial frequencies. eNeuro 6(5):ENEURO.0301-19.

Krakauer JW, Hadjiosif AM, Xu J, Wong AL, Haith AM. Motor learning. Compr. Physiol. 9, 613–663, 2019.

Kurtzer IL (2015) Long-latency reflexes account for limb biomechanics through several supraspinal pathways. Front Integr Neurosci 8:99.

Leow L-A, Gunn R, Marinovic W, Carroll TJ. Estimating the implicit component of visuomotor rotation learning by constraining movement preparation time. J Neurophysiol 118: 666–676, March 29, 2017.

Leow LA, Marinovic W, de Rugy A, Carroll TJ. (2020) Task errors drive memories that improve sensorimotor adaptation. J Neurosci 40:3075–3088.

Leow L-A, Marinovic W, de Rugy A, Carroll TJ. 2018. Task errors contribute to implicit aftereffects in sensorimotor adaptation. Eur. J. Neurosci 48 (11):3397–409.

Li S, Zhu Z, Adams S. An exploratory study of arm-reach reaction time and eye-hand coordination. Ergonomics38: 637–650, 1995.

Loeb, G. E. (2012). Optimal isn’t good enough. Biological cybernetics, 106, 757–765.

Mangin EN, Chen J, Lin J, Li N. (2023) Behavioral measurements of motor readiness in mice. Current Biology 33:3610–3624.

McPeek, R. M., Han, J. H., & Keller, E. L. (2003). Competition between saccade goals in the superior colliculus produces saccade curvature. J Neurophysiol, 89(5), 2577–2590

Meirhaeghe, N., Riehle, A., and Brochier, T. Parallel movement planning is achieved via an optimal preparatory state in motor cortex. Cell Reports, 42(2), 2023.

Morehead JR, Taylor JA, Parvin DE, Ivry RB. Characteristics of implicit sensorimotor Adaptation revealed by task-irrelevant clamped feedback. J Cogn Neurosci 29: 1061–1074, 2017.

Moullet E, Roby-Brami A, Guigon E. 2022. What is the nature of motor adaptation to dynamic perturbations? PLoS Comput Biol e1010470.

Munoz DP, Everling S. 2004. Look away: the anti-saccade task and the voluntary control of eye movement. Nat. Rev. Neurosci. 5, 218–228.

Nashed JY, Diamond JS, Gallivan JP, Wolpert DM, Flanagan JR. 2017. Grip force when reaching with target uncertainty provides evidence for motor optimization over averaging. Scientific Reports 7:11703.

Onagawa R, Kudo K (2021) Sensorimotor strategy selection under time constraints in the presence of two motor targets with different values. Sci Rep 11:22207.

Perich MG, Gallego JA, Miller LE. (2018) A neural population mechanism for rapid learning. Neuron 100(4):964–976.

Posner MI. Orienting of attention: then and now. Q J Exp Psychol (Hove) 69: 1864–1875, 2016.

Pruszynski, J. A. & Scott, S. H. Optimal feedback control and the long-latency stretch response. Exp. Brain Res. 218, 341–359 (2012).

Rao RP. A sensory–motor theory of the neocortex. Nature Neuroscience 1–15, 2024.

Raphael, G., Tsianos, G. A., & Loeb, G. E. (2010). Spinal-like regulator facilitates control of a two-degree-of-freedom wrist. Journal of Neuroscience, 30(28), 9431–9444.

Schween R, McDougle SD, Hegele M, Taylor JA. Assessing explicit strategies in force field adaptation. J Neurophysiol 123: 1552–1565, 2020.

Smeets JBJ, Wijdenes OL, Brenner E (2016) Movement adjustments have short latencies because there is no need to detect anything. Mot Control 20:137–148.

Steinmetz, N. A., Zatka-Haas, P., Carandini, M. & Harris, K. D. Distributed coding of choice, action and engagement across the mouse brain. Nature 576, 266–273 (2019).

Stewart BM, Baugh LA, Gallivan JP, Flanagan JR. 2013. Simultaneous encoding of the direction and orientation of potential targets during reach planning: evidence of multiple competing reach plans. Journal of Neurophysiology 110:807–816. DOI: 10.1152/jn.00131.2013, PMID: 23699052

Stewart BM, Gallivan JP, Baugh LA, Flanagan JR. 2014. Motor, not visual, encoding of potential reach targets. Current Biology 24:R953–R954.

Taylor JA, Krakauer JW, Ivry RB. Explicit and implicit contributions to learning in a sensorimotor adaptation task. J Neurosci 34: 3023–3032, 2014.

Tsianos, G. A., Goodner, J., & Loeb, G. E. (2014). Useful properties of spinal circuits for learning and performing planar reaches. Journal of neural engineering, 11(5), 056006.

Van der Stigchel S, Meeter M, Theeuwes J. 2006. Eye movement trajectories and what they tell us. Neuroscience & Biobehavioral Reviews 30:666–679.

Veerman MM, Brenner E, Smeets JBJ (2008) The latency for correcting a movement depends on the visual attribute that defines the target. Exp Brain Res 187:219–228.

Watanabe R, Higuchi T. Anticipatory action planning for stepping onto competing potential targets. Front. Hum. Neurosci. 2022, 16, 875249.

Weiler J, Gribble PL, Pruszynski JA (2019) Spinal stretch reflexes support efficient hand control. Nat Neurosci 22:529 –533.

Weiler J, Gribble PL, Pruszynski JA (2021) Spinal stretch reflexes support efficient control of reaching. J Neurophysiol 125:1339–1347.

Wijdenes OL, Brenner E, Smeets JBJ (2014) Analysis of methods to determine the latency of online movement adjustments. Behav Res Methods 46(1):131–139.

Wong AL, Goldsmith J, Forrence AD, Haith AM, Krakauer JW (2017) Reaction times can reflect habits rather than computations. eLife 6e28075.

Wong AL, Green AL, Isaacs MW (2022) Motor plans under uncertainty reflect a trade-off between maximizing reward and success. eNeuro 9:ENEURO.0503-21.2022.

Wong AL, Haith AM, Krakauer JW. Motor planning. Neuroscientist 21: 385–398, 2015.

Wong AL, Haith AM. 2017. Motor planning flexibly optimizes performance under uncertainty about task goals. Nature Communications 8:14624.

Zhang Y, Brenner E, Duysens J, Verschueren S, Smeets JBJ (2018a) Effects of aging on postural responses to visual perturbations during fast pointing. Front Aging Neurosci 10:401.

Zhang Y, Brenner E, Duysens J, Verschueren S, Smeets JBJ (2018b) Postural responses to target jumps and background motion in a fast pointing task. Exp Brain Res 236:1573–1581.

Zhou W, Kruse EA, Brower R, North R, Joiner WM. (2022). Motion state-dependent motor learning based on explicit visual feedback is quickly recalled, but is less stable than adaptation to physical perturbations. J. Neurophysiol. 128, 854–871.

